# Biphasic growth dynamics during *Caulobacter crescentus* division

**DOI:** 10.1101/047589

**Authors:** Shiladitya Banerjee, Klevin Lo, Matthew K. Daddysman, Alan Selewa, Thomas Kuntz, Aaron R. Dinner, Norbert F. Scherer

## Abstract

Cell size is specific to each species and impacts their ability to function. While various phenomenological models for cell size regulation have been proposed, recent work in bacteria have demonstrated an *adder* model, in which a cell increments its size by a constant amount between each division. However, the coupling between cell size, shape and constriction, remain poorly understood. Here, we investigate size control and the cell cycle dependence of bacterial growth, using multigenerational cell growth and shape data for single *Caulobacter crescentus* cells. Our analysis reveals a biphasic mode of growth: *a relative timer* phase before constriction where cell growth is correlated to its initial size, followed by a *pure adder* phase during constriction. Cell wall labeling measurements reinforce this biphasic model: a crossover from uniform lateral growth to localized septal growth is observed. We present a mathematical model that quantitatively explains this biphasic *mixer* model for cell size control.

---

We recently introduced a technology that enables obtaining unprecedented amounts of precise quantitative information about the shapes of single bacteria as they grow and divide under non-crowding and controllable environmental conditions [1, 2]. Others have developed complementary methods [3–6]. These single-cell studies are generating great interest because they reveal unanticipated relationships between cell size and division control [5]. Recent work in bacteria revealed a model of constant size increment between successive generations for a wide range of bacterial species [3–5, 7, 8], as originally proposed in Ref. [9], and recently termed as an *adder* model [5, 10]. Competing models for size control include cell division close to a critical size (*sizer*) [11] or at a constant interdivision time (*timer*), equivalent to a critical multiple of the birth size with a constant growth rate [1]. Analysis of single-cell data show that cell size at division is positively correlated with the cell size at birth [1, 4, 5, 12, 13], thus precluding a sizer model. In addition, a negative correlation between initial cell size and interdivision times, as reported here and in refs [1, 4, 5, 13, 14], is inconsistent with the timer model. However, other studies have suggested mixed models of size control, with diverse combinations of sizer, timer and adder models [10, 15–17]. The spatial resolution and statistically large size of our data now allow us to revisit these issues with greater precision.

While cell size serves as an important determinant of growth, the bacterial cell cycle is composed of various coupled processes including DNA replication and cell wall constriction that have to be faithfully coordinated for cells to successfully divide [18]. This raises the question of what other cell cycle variables regulate growth and how the interplay between these variables can be understood quantitatively [19, 20]. Indeed, our recent modeling and analysis of cell shape dynamics revealed how different shape parameters are coupled through growth and division [2, 21]. Here we relate cell size control and cell wall growth to the timing of cell-wall constriction in *C. crescentus* cells.

## RESULTS

We use a combination of microfluidics and phase-contrast microscopy for high-throughput, live-cell measurements of cell shape dynamics of single *C. crescentus* cells [1, 2, 22]. As a population of cells is controllably attached via a stalk and holdfast to the coverslip surface in the microfluidic channel, our measurements allow obtaining accurate and precise data of single cell shape and growth for >10000 generations for >250 cells under steady environmental conditions. From the splined cell contours of the acquired phase-contrast images (Fig. 1a), we determine various cell shape parameters, such as the length of the cell midline axis (*l*), cell width, and the radius of curvature of the midline. As reported previously, *l* increases exponentially, *l*(*t*) = *l*(0)*e^κt^*, with time constant 〈*κ*〉^−1^ = 125 ± 8 min and a mean interdivision time 〈*τ* 〉 = 73 ± 7 min at 31°C in peptone-yeast extract (PYE) medium, while the average width and the radius of curvature remains approximately constant [2]. Since measurements of the cell area behave the same as the length [1], we use the length as a metric for cell size.

**FIG. 1.**
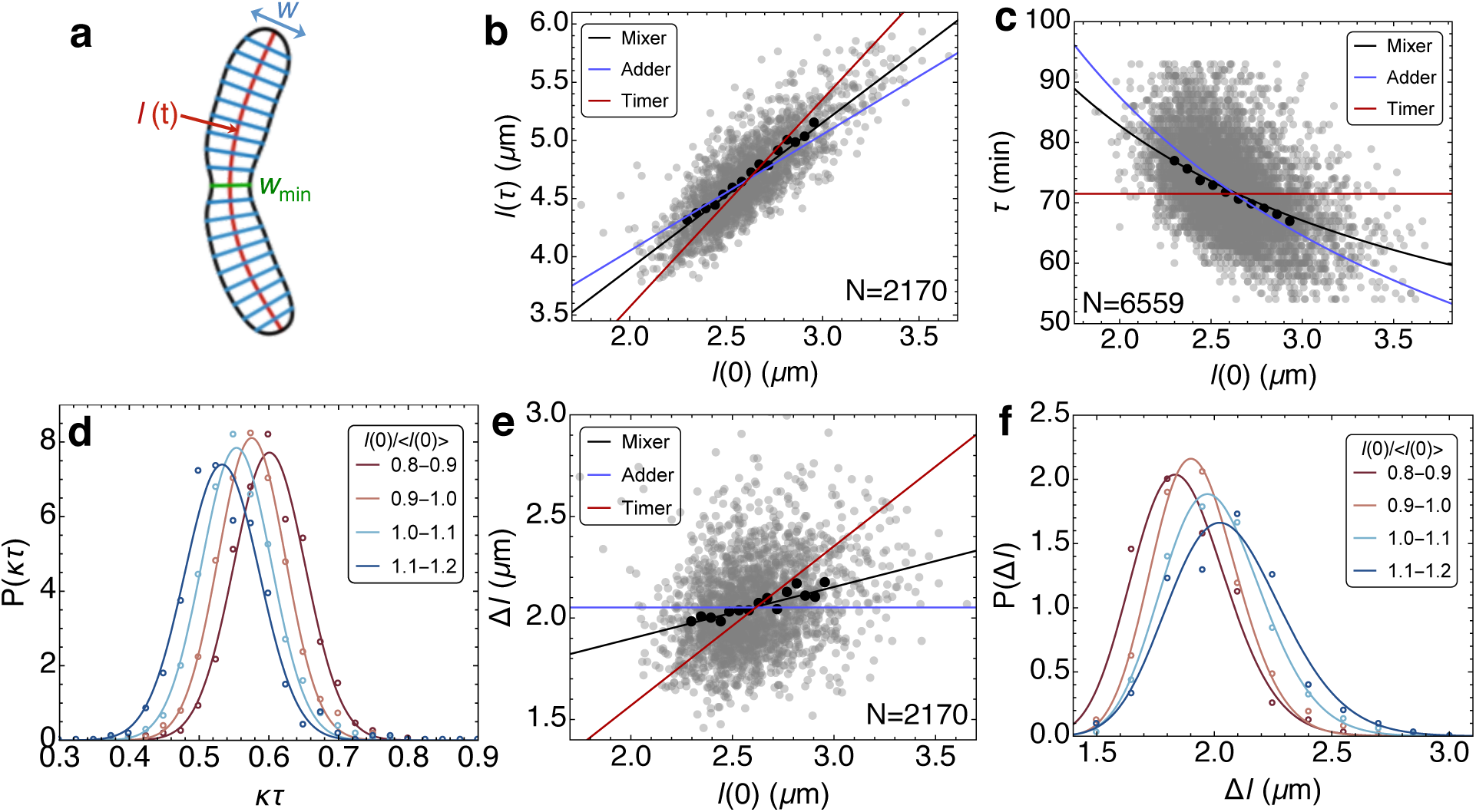
Cell size and division control in *C. crescentus*. **a.** A representative splined contour of a *C. crescentus* cell, illustrating the shape variables. **b.** Cell size at division, *l*(*τ*), vs the cell size at birth, *l*(0). The black solid line represents a least square linear fit to all single generation data given by the gray scatter cloud i.e. mixer model. Corresponding fits by timer and adder models are given by red and blue lines, respectively. The solid circles represent mean data binned by *l*(0). **c.** The scatter plot of interdivision times, *τ*, vs the initial cell size, *l*(0), exhibits a negative correlation. The black solid line is the prediction based on exponential growth and the mixer model with no adjustable fitting parameters. Predictions from timer and adder models are shown by the red and blue lines, respectively. **d.** The conditional probability density of normalized division cycle time, *κτ*, given the mean rescaled initial length values, *P* (*κτ* |*l*(0)*/*〈*l*(0)〉), illustrating the negative feedback between *τ* and *l*(0). The open circles represent experimental data and the solid curves are Gaussian fits. **e.** Size extension in each generation, Δ*l* is correlated with the initial cell size. The mean trend is described by the linear relationship, Δ*l* = (*a* − 1)*l*(0) + *δ*, which is the mixer model. The solid circles represent mean data binned by *l*(0). **f.** shows the conditional probability density of size extension Δ*l* given the mean rescaled initial cell length, *P* (Δ*l*|*l*(0)*/*〈*l*(0)〉). The open circles represent experimental data and the solid curves are lognormal fits. *N* indicates number of generations in **b**, **c**, **e**.

### Mixer model of cell size control

We first analyzed the correlation between cell size at birth, *l*(0), and at division, *l*(*τ*), which describes the strategy for cell size control. Previously [1], the relationship between cell size at birth and at division was described by fitting the data with only pure timer (*l*(*τ*) = *al*(0), *a* is a proportionality constant) and adder (*l*(*τ*) = *l*(0) + *δ*) models. Here, consistent with [10], we find that cell size correlation in that same dataset can be best described by a model that combines both adder and timer components: *l*(*τ*) = *al*(0) + *δ*, with a slope of *a* = 1.25 and an intercept *δ* = 1.39 *μ*m (Fig. 1b; Supplementary Note 1; Supplementary Fig. 1). The value of the slope should be contrasted with 1.8 [1], the multiple expected for a size ratio ≃0.55 between the daughter cells. While the interdivision times, *τ*, and the growth rates, *κ*, fluctuate between cells and across generations, positive *δ* implies that larger cells divide more quickly than smaller cells [4, 5, 7, 13],
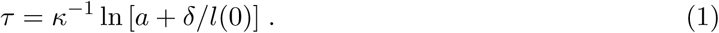

We find *τ* to be negatively correlated with *l*(0) (Eq. (1); Fig. 1c) [4]. As shown in Fig. 1d, the distributions of normalized division cycle times, *κτ*, are also correlated with the initial lengths as shown by the conditional probability *P*(*κτ*|*l*(0)*/*〈*l*(0)〉) for various ranges of *l*(0)*/*〈*l*(0)〉. These observations rule out a timer model of size control where the division times would be uncorrelated with the initial lengths [1]. Furthermore, Fig. 1e-f show that the lengths added in each generation, Δ*l* = *l*(*τ*) − *l*(0), are positively correlated with the initial cell lengths, which precludes a pure adder model for cell size control, in contrast to [4]. Our data suggest that *C. crescentus* cells behave with attributes of both timer and adder, i.e. a *mixer*. Furthermore, we find that this mixer model is conserved in the temperature range 14°C-34°C (Supplementary Fig. 2, 3). Interestingly, cells at 37°C behave as a perfect adder, which suggests that adder-like behavior may be elicited by experimental conditions. For *C. crescentus*, 37°C is the extreme upper limit for viable cell growth.

**FIG. 2.**
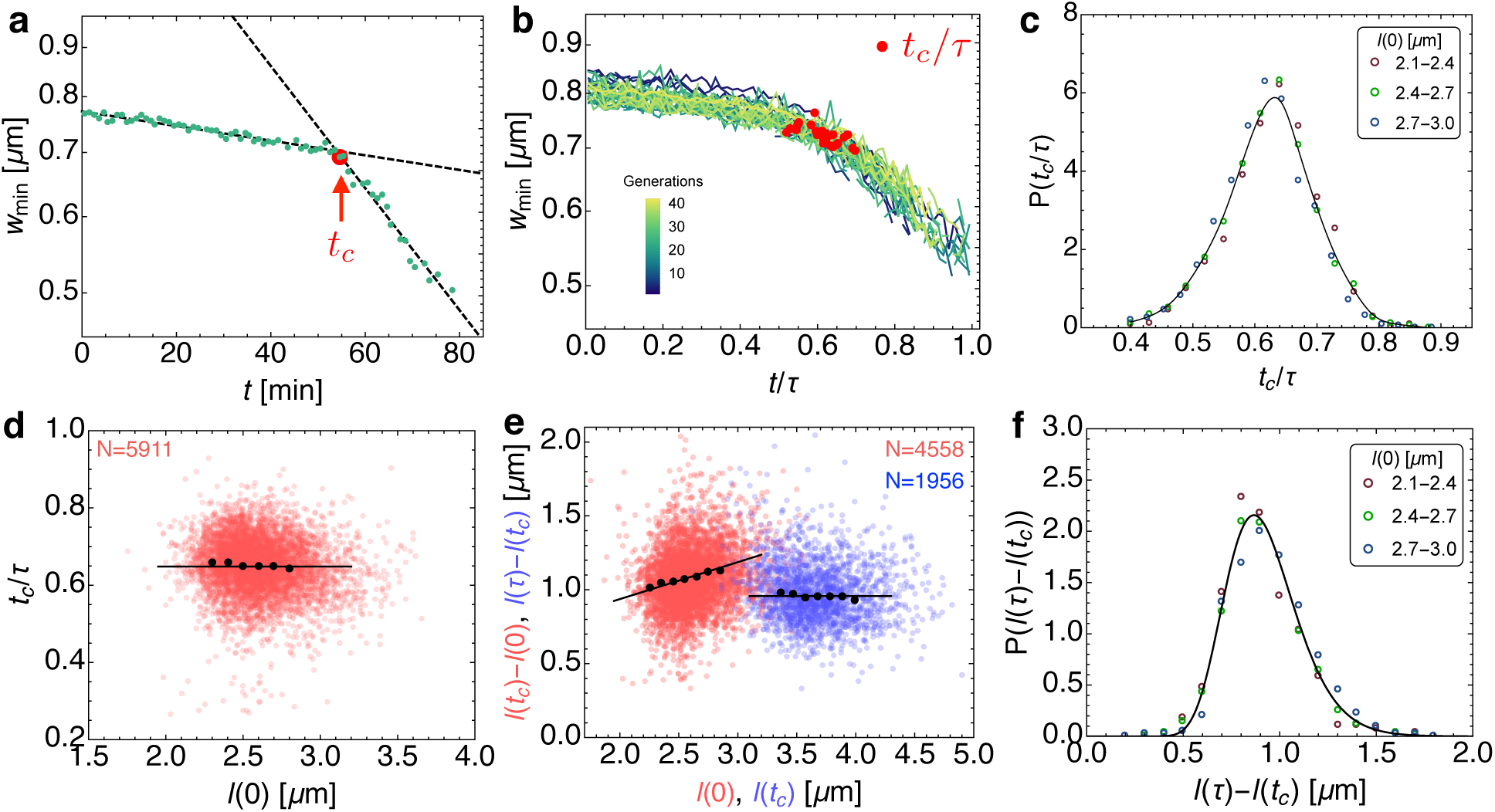
Crossover from relative timer to adder at the onset of cell wall constriction. **a.** Semi-log plot of the time-dependence of *w*_min_ in a representative generation shows two phases of constriction, a slow initial phase followed by a fast constriction phase. We determine *t_c_* by the intersection of the least square exponential fits to the earlier and later portions of the division cycle (Methods). **b.** Dynamics of *w*_min_ across generations of a typical cell as functions of the normalized division cycle time. Locations of *t_c_/τ* are marked by red solid circles. **c.** The conditional probability density of the normalized crossover time, *t_c_/τ*, given initial length values, *P* (*t_c_/τ* |*l*(0)), shown by colored circles. The density indicates that *t_c_/τ* and *l*(0) are independent. Solid line is a best fit cubic spline curve. **d.** Red scatter points and mean values (black points) show a lack of correlation between *t_c_/τ* and *l*(0). Black line represents a relative timer: *t_c_* = 0.63*τ*. The solid circles represent mean data binned in *l*(0). **e.** Positive correlation between the added size before constriction, *l*(*t_c_*) − *l*(0) and *l*(0) (red scatter). The black line represents the best fit: *l*(*t_c_*) = 1.25*l*(0) + 0.43. Added size for *t > t_c_*, is uncorrelated with *l*(*t_c_*) (blue scatter) supporting a pure adder model during the constriction phase: *l*(*τ*) = *l*(*t_c_*) + 0.97. **f.** The conditional probability density of the post-constriction added cell size. Colored circles indicate the ranges of the initial lengths, *l*(0). The collapse of the distributions indicates the independence of *l*(*τ*) − *l*(*t_c_*) and *l*(0). The solid line is a best-fit lognormal distribution. *N* indicates number of generations in **d** and **e**.

### Relative timer phase prior to cell wall constriction

Since a mixer model implements a timer and an adder component serially, we examined our single cell shape data [2] to determine a crossover in growth behavior. We find that the constriction dynamics in individual generations exhibit a biphasic behavior, with an initial period of slow constriction followed by a phase of fast constriction (Fig. 2a). We determine the crossover time, *t_c_*, by fitting piecewise exponential curves to the initial and the later phases of decay in *w*_min_(*t*) (Methods; Supplementary Fig. 5 a-b). We estimate the onset of constriction by *t_c_*, which has a mean value *t_c_* = 47 ± 7 min at 31°C (Supplementary Fig. 5c). The data for *w*_min_ across cell lineages collapse to a master crossover curve when time is normalized by interdivision times (Fig. 2b-c), indicating that a single timescale governs constriction initiation. This crossover dynamic is observed in the analogous data obtained at other temperatures of the medium (Supplementary Fig. 6). We find that *t_c_* increases in proportion to *τ* and *κ*^−1^ as the temperature is decreased (Supplementary Fig. 7). The conditional distributions of the normalized crossover times, *t_c_/τ*, shown in Fig. 2c, collapse to a single curve for various values of *l*(0), independent of initial cell length. Indeed our data show that *t_c_/τ* is nearly uncorrelated with the initial cell size (Fig. 2d), whereas the cell length at *t* = *t_c_* increases in proportion to the initial length, *l*(*t_c_*) = 1.25 *l*(0) + 0.43 (Fig. 2e). By analyzing our shape data at other temperatures we find that *t_c_/τ* remains independent of *l*(0) (Supplementary Fig. 9a), and does not vary with changing temperature of the growth medium (Supplementary Fig. 9b).

### Pure adder phase during cell wall constriction

While the time to the onset of cell wall constriction is uncorrelated with cell size, the added size in the constriction phase, *δ*^′^ = *l*(*τ*) − *l*(*t_c_*), also shows no correlation with *l*(*t_c_*) (Fig. 2e). This suggests a *pure adder* model of cell size control for *t_c_ < t < τ*, such that the distribution of the added size is independent of the initial cell length. This adder behavior is confirmed by the collapse of the conditional distributions of the added size *P* (*δ*^′^|*l*(0)〉) to a single curve, approximated by a log-normal distribution (Fig. 2f) [7]. Furthermore, the time to divide after *t_c_* shows negative correlation with *l*(*t_c_*) (Supplementary Fig. 8). This negative correlation is supported by an *adder* model for *t > t_c_*, *τ* = *t_c_* + *κ*^−1^ ln [1 + *δ*^′^*/l*(*t_c_*)] with *δ*^′^ = 0.97 *μ*m. We find that the adder phase post constriction is conserved for all temperatures with a mean added size ≈ 1 *μ*m (Supplementary Fig. 9 c-d).

### Crossover in cell wall growth dynamics

We conducted fluorescence labeling experiments to determine if we can visualize a crossover from timer to adder growth phase by selective labeling. We sought to examine the division cycle dependence of peptidoglycan synthesis, by using a fluorescent construct of lectin wheat germ agglutinin (flWGA) that has been shown to label the cell wall of Gram-negative bacteria [23]. Consistently, our experiments measuring the WGA fluorescence of stalked *Caulobacter* cells showed peripheral WGA localization in confocal slices (Supplementary Fig. 10). Using our microfluidics platform [1], *C. crescentus* cells were initially incubated in media with PYE and flWGA for 15 minutes without flow, allowing the cells to be covered with flWGA. PYE media was then flowed into the microfludics channel and image stacks of stalked cells were acquired every 10 minutes within the fields of view. The deconvolved images in Fig. 3a show that the flWGA intensity is spatially uniform prior to constriction (i.e., for samples at *t* < 50 min), but exhibits a pronounced minimum at the septum as the cell-wall is invaginated (*t* > 50 min). Moreover, the 70 and 80 min images even hint at the secondary invaginations in a predivisional cell, consistent with our previous report [2]. For each of these images, Fig. 3b shows the intensity along the centerline axis, averaged over the cell cross-section at each position and then normalized by the maximum value for each time. Because we account for variation of the cell cross-section, the appearance of the minimum in the intensity at the septum is not an artefact of its diminishing width. The spatial distribution of flWGA intensity suggests that growth is spatially uniform for *t* < 50 min and new cell-wall material is primarily synthesized at the invagination for *t* > 50 min. This septal mode of growth has been reported earlier with D-amino acid cell wall labeling [24, 25].

**FIG. 3.**
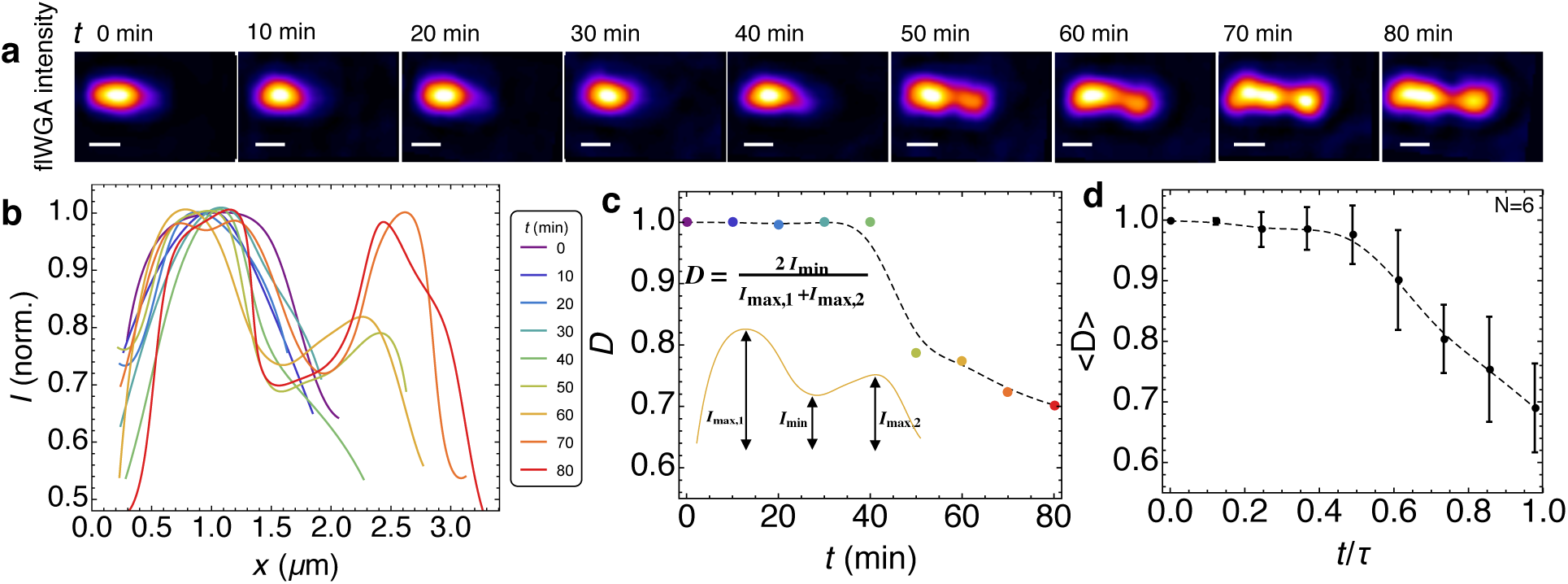
Crossover in cell wall growth dynamics at the onset of constriction. **a.** Confocal fluorescent images of a representative *C. crescentus* cell in a microfluidic flow cell labeled with fluorescent WGA taken after 0, 10, 20, 30, 40, 50, 60, 70 and 80 min of growth in PYE medium. The fluorescence intensity is averaged over cell thickness (see Supplementary Fig. 10 for mid-plane fluorescence). The images are deconvoluted using the Huygens software package (Methods). The scale bars represent 1 *μ*m. The depletion of fluorescence reveals the underlying spatial pattern of growth, i.e. growth occurs where the fluorescence is minimized. **b.** Spatial distribution of flWGA intensity along the centerline axis, averaged over the cell cross-section at each position. We then normalized by the maximum value for each time to account for variations in [$]WGA labeling. The cross-section averaging accounts for the change in surface area reduction in the septal region. The time points are indicated with colors progressing from purple to red. **c.** Inset: A typical intensity profile is characterized by one minimum at the septum (*I*_min_) and two maxima near either pole (*I*_max,1_, *I*_max,2_). We define the index of uniformity as *D* = 2*I*_min_/(*I*_max,1_ + *I*_max,2_) (Methods). *D*(*t*) is shown for a representative cell in (A), revealing a crossover from uniform growth (〈*D*〉 ≃ 1) to localized septal growth at *t* ~ 50 min. **d.** Ensemble averaged dynamics of the growth uniformity index, 〈*D*〉, as a function of time normalized by the division time. Error bars indicate ±1 standard deviation.

To quantify the spatial uniformity of cell-wall deposition for each cell in the ensemble we introduce an *intensity uniformity index*, *D*, given by the ratio of the intensity at the site of the septum (*I*_min_) to the mean of the maximum intensities (*I*_max,1,2_) on the stalked and swarmer sides of that site (Fig. 3c-inset). *D* is close to unity for *t* < 50 min (since *I*_min_ ≃ *I*_max,1,2_), indicating spatially uniform growth (Fig. 3c). For *t* > 50 min, *D* drops sharply to lower values, suggesting that cell wall growth is localized to the septum. Fig. 3d shows the ensemble averaged intensity uniformity index, 〈*D*〉, exhibiting a smooth crossover to septal growth for *t* > 0.6*τ*.

### Septal growth model describes biphasic constriction

To examine whether septal cell wall synthesis (Fig. 3) can reproduce the observed crossover dynamics of constriction (Fig. 2), we consider a quantitative model for cell wall constriction driven by septal growth [21, 26]. We assume that the shape of the constriction zone is given by two intersecting hemispherical segments with diameter *w*, and constriction proceeds by completing the missing parts of the hemispheres while maintaining the curvature of the preformed spherical segments. The total surface area of the septum is given by *S*(*t*) = *πwl_s_*(*t*), where *l_s_*(*t*) is the total length of the hemispherical segments (Fig. 4a). Exponential growth of septal surface area implies [2]:
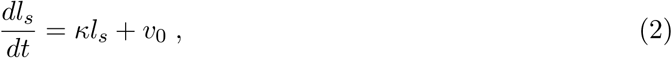

where *v*_0_ is the speed of septum synthesis at *t* = 0, which we determine by fitting our model to the data for *w*_min_(*t*). Eq. (2) can be solved using the initial condition, *l_s_*(0) = 0, to derive the time-dependence of *w*_min_(*t*),

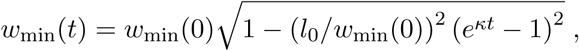

where *l*_0_ = *v*_0_*κ*^−1^. As a result, the dynamics of constriction are controlled by the dimensionless parameter, aspect ratio *l*_0_/*w*_min_(0), exhibiting a crossover point (Fig. 4b). As shown in Fig. 4b, the model prediction for the crossover time is in excellent agreement with our experimental data, and the time to the onset of constriction is insensitive to variations in *l*_0_/*w*_min_(0) (Supplementary Note 2; Fig. 4b-inset). This implies that *t_c_* is controlled by *κ*, which is consistent with the positive correlation between *t_c_* and *κ*^−1^ (Fig. 4c). The septal growth model describes a smooth transition from predominantly lateral growth to predominantly septal growth, consistent with the data. We explicitly show how the dynamics depend on the rate of transition in Supplementary Note 3 (Supplementary Fig. 12).

**FIG. 4.**
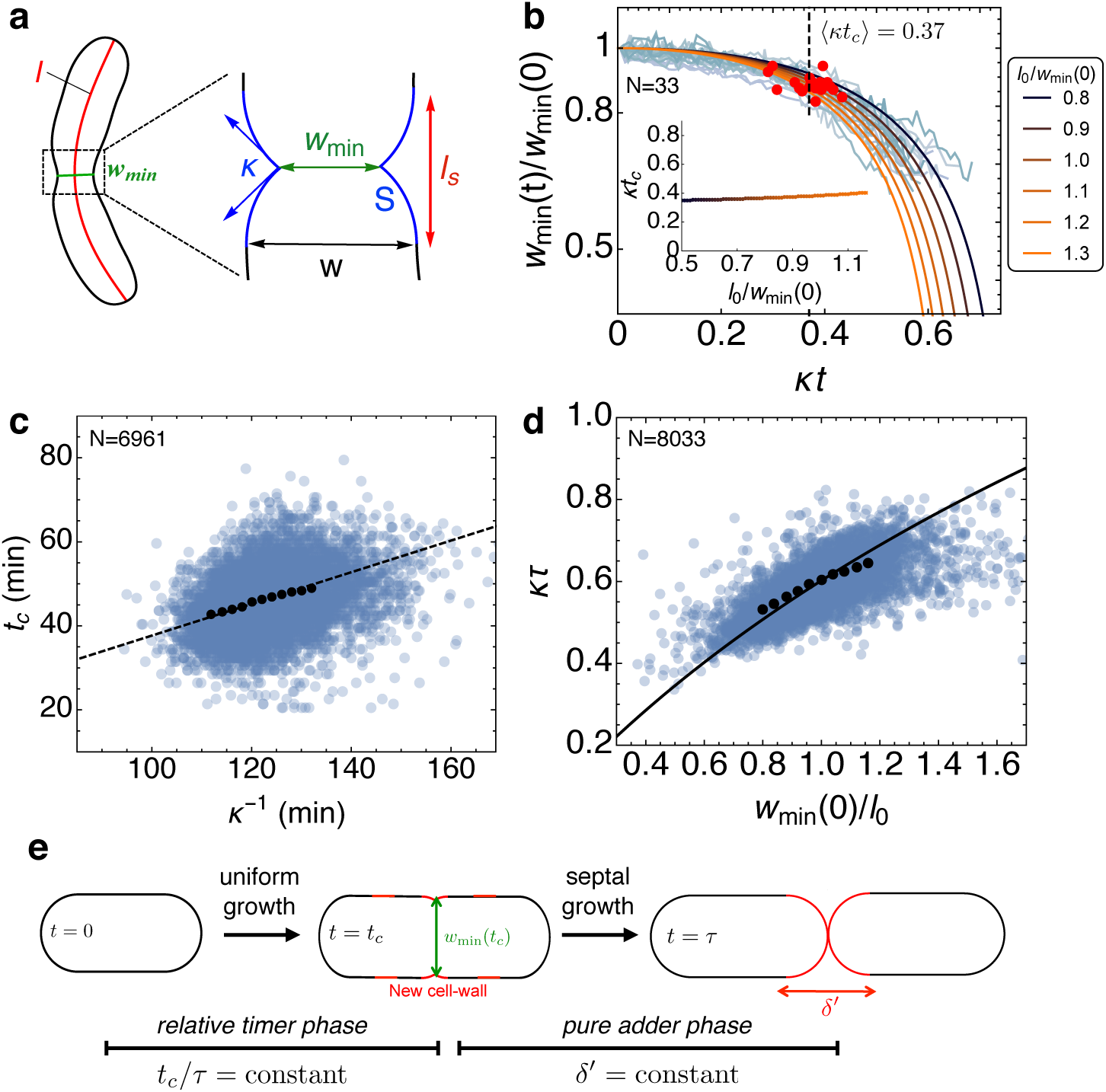
Septal growth model predicts the onset of cell wall constriction and interdivision times. **a.** A representative splined contour of a *C. crescentus* cell, illustrating the shape parameters, *l* (red) and *w*_min_ (green). The region inside the dashed rectangle represents the constriction zone, where *S* is the surface area of the septal cell-wall (blue) synthesized after constriction. **b.** Dynamics of the width of the pinch-off plane (normalized by *w*_min_(0)), with time normalized by *κ*^−1^. Predictions of the septal growth model are shown by solid curves at various values of the dimensionless parameter, *l*_0_/*w*_min_(0). Experimental data for different generations of a representative cell are shown in light blue with the locations of the crossover marked by solid red circles. Inset: No dependence of *κt_c_* on *l*_0_/*w*_min_(0), as predicted by the theoretical model (Supplementary Note 2). **c.** Positive correlation between *t_c_* and *κ*^−1^ (blue scatter). The solid black circles represent mean data binned in *κ*^−1^ and the dashed line represents the best fit with *t_c_* = 0.37*κ*^−1^. **d.** Normalized interdivision times (*κτ*) vs the normalized initial width of the pinch-off plane (binned mean data as solid black circles; model prediction as solid curve; see Supplementary Note 4; Supplementary Fig.13a). **e.** Schematic for spatiotemporal coordination of cell wall growth in *C. crescentus* cells. Growth is spatially uniform for *t* < *t_c_* when cell wall deposition occurs along the entire cell length. For *t > t_c_*, cell wall growth is dominated by septal cell wall synthesis that leads to constant size extension determined by the surface area of the new poles. *N* indicates number of generations in **b**, **c** and **d**.

Another prediction of the septal growth model is that the interdivision times increase with *w*_min_(0)/*l*_0_, because wider cells require more material to close the septum. Based on our model, we predict a simple relation between *τ* and *w*_min_(0) (Supplementary Note 4),
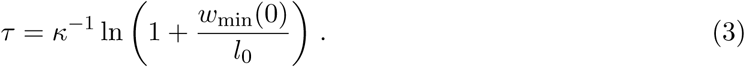

Thus, the interdivision time is predicted to be longer for larger *w*_min_(0). We find a positive correlation in our data between *κτ* and *w*_min_(0)/*l*_0_, and the mean trend in our data is in good quantitative agreement with our model prediction (Fig. 4d). Since constricting cells primarily grow from the septum (Fig. 3), the added size in the constriction phase, *δ*^′^, is expected to be proportional to the width of the septal plane. We find that *δ*^′^ is positively correlated with *w*_min_(*t_c_*) (Supplementary Fig. 13b). Furthermore, Eqs. (1) and (3) together imply a negative correlation between initial septal width, *w*_min_(0), and the cell size at division, *l*(*τ*), in quantitative agreement with our data (Supplementary Note 5; Supplementary Fig. 13c).

## DISCUSSION

The adder phase of cell-wall growth (*t > t_c_*) coupled with relative timer prior to constriction (*t < t_c_*) comprise a biphasic growth model for *C. crescentus* (Fig. 4e). Newborn cells exhibit uniform patterning of cell wall synthesis prior to constriction (*t < t_c_*), and initiate constriction at a fixed phase in the division cycle. During the constriction phase (*t > t_c_*), cells primarily add new cell wall material at the septum. This phase of growth is characterized by a constant cell size extension, proportional to the cell width. The length added after *t > t_c_* originates primarily from the surface area of the daughter cell poles. Taken together, the crossover from uniform cell wall growth to localized septal growth provides a physical basis for the *mixer model* of cell size control.

Like the adder model, the mixer model ensures cell size homeostasis: with each division, the cell length regresses to the ensemble average [10]. It is interesting to consider why *E. coli* exhibits a pure adder behavior while *C. crescentus* exhibits a mixer behavior. One notable difference between the species is that *E. coli* can initiate multiple rounds of DNA replication per cell cycle while *C. crescentus* has a strict one-to-one correspondence between these processes. While the molecular basis for the biphasic cell size control is unknown, the relative timer phase may be related to the duration of chromosome replication, which is independent of cell size. Alternatively, it has been suggested that nutrient uptake imposes condition-dependent constraints on surface-area-to-volume ratios and in turn the growth mechanism [17]. Our study provides additional insights for investigating the molecular candidates regulating cell size and division control in bacteria.

## METHODS

### Acquisition of Experimental Data and Cell Shape Analysis

Experimental data were acquired as described in [1]. In the main article we use the exact same dataset as in refs. [1, 2], consisting of 260 cells, corresponding to 9672 generations (division events) at 31°C. Corresponding data and analysis of cell shape for other temperature are provided in the Supplementary Figures. The acquired phase-contrast images were analyzed using a custom routine in Python [1, 2]. See Supplementary Methods for further details.

### WGA Fluorescence Microfluidics Assay

To investigate the dynamics of cell wall growth over time in *Caulobacter crescentus*, we monitored the localization of fluorescent wheat germ agglutinin (flWGA) on the cell wall of the bacteria using the microfluidics platform we previously developed [1]. A 5 mL liquid culture of *C. crescentus* in PYE was prepared overnight and diluted the following morning to an optical density at 660 nm of 0.1. Vanillate was added to this diluted culture at a final concentration of 0.5 mM to induce the production of the holdfast hfsA for 3 hrs. 1 mL of this culture was flowed into a cleaned microfluidics chip and allowed to incubate for 1 hr. A 20 mL syringe of PYE and a 3 mL syringe with 2 mL of PYE and 1 mL of flWGA were attached to two separate input ports into the microfluidics channel. Flow into the channel was resumed with media from the 3 mL syringe at a rate of 3.5 L/min for 15 minutes. Flow was then halted for 15 minutes to allow the cells to be covered with the flWGA. Media from the 20 mL syringe was flowed into the channel at a rate of 3.5 L/min. Image stacks with a 100 nm spacing were acquired every 10 minutes (using MicroManager) at a position along the microfluidics channel with sufficient cell coverage.

### Deconvolution of Images

Captured images of an object (*I*) are a convolution of the actual 11 object (*f*) and the point spread function (PSF) of the microscope (*h*):

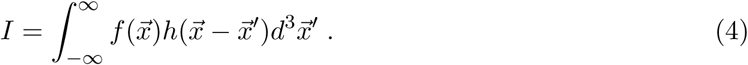

Given a measurement of the PSF and the acquired images of the object, our deconvolution approach employs a classic maximum-likelihood estimation algorithm that calculates the most likely object to produce the acquired images [27]. This calculation is performed relatively quickly in the Fourier domain, where the integral (4) is transformed into simple multiplication. Deconvolution microscopy is widely used to remove the blurring imposed by the PSF of the microscope.

An image stack from each time point was deconvolved individually using commercial software (Huygens Deconvolution; Scientific Volume Imaging). Before performing the deconvolution, the 3D PSF of our oil immersion objective (Nikon), with a magnification of 100X and NA = 1.49, was measured by imaging static 100-nm-diameter polystyrene beads coated with green fluorescent protein (Thermofisher). The PSF was sampled at 72 nm by 72 nm in the x-y plane and 50 nm in the z direction, thus satisfying the Nyquist criteria for our particular objective. Next, 17 image stacks corresponding to 17 time points were loaded into Huygens. Parameters such as background intensity, spatial sampling, objective NA, immersion index of refraction, and the signal-to-noise ratio (SNR) of objects were entered manually.

The background intensity was determined by calculating the mean intensity in an area of the image where there is no signal. The bacterial image stacks were sampled at 72 nm by 72 nm in the x-y plane and 100 nm in the z-direction. Because of photo-bleaching, the SNR of the bacteria will change as a function of time. At each time point, the SNR was calculated using the following equation: 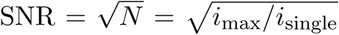, where *i*_max_ is the maximum grayscale value of a pixel in the bacteria and *i*_single_ is the grayscale value due to a single photon incident on our detector. The value of *i*_mean_ is obtained at each time point from the measured images. The value of *i*_single_ is a calculated quantity using the parameters of our camera (Andor iXon EMCCD) such as quantum efficiency, A/D conversion, and system gain. A maximum number of 40 iterations was allowed for the deconvolution, but Huygens reached a global minimum at ~30 iterations for each time point.

### Intensity uniformity index

A typical intensity profile at early times (*t* < 50 min) is spatially uniform around the cell center and then decays towards the poles. At later times, *t* > 50 min, the intensity profile is characterized by one minimum at the septum, given by *I*_min_, and two maxima at the stalked and the swarmer components, given by *I*_max,1_ and *I*_max,2_ respectively (Fig. 3c - inset). At each time point, we define the growth uniformity index for each intensity profile as, *D* = 2*I*_min_/(*I*_max,1_ + *I*_max,2_). *I*_min_ is defined as the minimum in the intensity profile for *r* − 2*σ < x*/*l < r* + 2*σ*, where *x* is the coordinate along the centerline, *r* is the mean ratio of the daughter cell lengths, and *σ* is the standard deviation in daughter cell length ratio. Let *x*_min_ denote the location of *I*_min_ along the centerline coordinate. Then *I*_max,1_ is defined as the maximum in the intensity for *x* < *x*_min_ and *I*_max,2_ is the maximum in the intensity profile for *x* > *x*_min_. Thus, for *t* ≤ 50, *I*_min_ ≃ *I*_max,I_ ≃ *I*_max,2_ and *D* ≃ 1. Whereas for *t* ≥ 50 min, *I*_min_ represents the flWGA intensity value at the septum and is lower than both *I*_max,1_ and *I*_max,2_.

### Crossover analysis from experimental data

To determine the crossover time, *t_c_*, from the data on *w*_min_, we fit the following piecewise linear function to ln (*w*_min_):

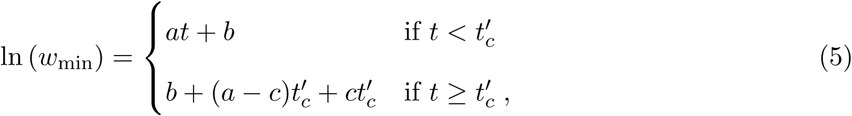

with four undetermined parameters *a*, *b*, *c* and 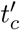 obtained using a built-in curve fitting function in Mathematica. A representative fit is given in Supplementary Fig. 5a, where 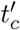 is the point of intersection of the two lines. We then compute the metric 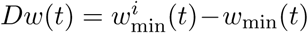, where 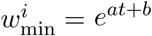 (Supplementary Fig. 5b). Constriction is estimated to initiate when the metric *Dw* exceeds a threshold of 0.05 *μ*m, which is equal to a single image pixel. The crossover time, *t_c_*, is taken to be 3 frames prior to the frame when *Dw* crosses the threshold value (Supplementary Fig. 5b), such that the determination of *t_c_* is robust to noise. We find the location for *t_c_* is significantly spread out across generations when *w*_min_(*t*) is plotted against absolute time (Supplementary Fig. 5c). When the constriction curves are aligned from the end of the cycle, as in Ref. [17], the individual curves collapsed, although the spread for *t_c_* is significant (Supplementary Fig. 5d). By contrast, when the constriction curves are plotted against relative time, as shown in Fig. 2b, the locations of the crossover, *t_c_/τ*, are much better aligned across generations.

### Data availability

The data that support the findings of this study are available from the corresponding authors upon request.

## ACKNOWLEDGEMENTS

We thank Charles Wright and Srividya Iyer-Biswas for measurements and shape analysis of *C. crescentus* single cell data [1, 2]. We thank Sean Crosson and Aretha Fiebig for contributing reagents, materials and helpful discussions. We gratefully acknowledge funding from the National Science Foundation Physics of Living Systems (NSF PHY-1305542), the National Science Foundation Materials Research Science and Engineering Center (MRSEC) at the University of Chicago (NSF DMR-0828854), the W. M. Keck Foundation and the Graduate Program in Biophysical Sciences at the University of Chicago (T32 EB009412/EB/NIBIB NIH HHS/United States). SB acknowledges support from the University College London for completion of part of this work.

## AUTHOR CONTRIBUTIONS

S.B., K.L., A.R.D. and N.F.S. designed research; S.B., K.L., A.S., M.D. and T.K. performed research; S.B., A.R.D. and N.F.S. wrote the manuscript.

